# ACE2 decoy Fc-fusions and bi-specific killer engager (BiKEs) require Fc engagement for *in vivo* efficacy against SARS-CoV-2

**DOI:** 10.1101/2024.06.20.599956

**Authors:** Jenna K. Dick, Dustin Hicks, Venkatramana D Krishna, Jules A. Sangala, Benjamin T. Zandstra, Carly Baehr, J Sjef Verbeek, Mark S. Cragg, Maxim C-J Cheeran, Marco Pravetoni, Geoffrey T Hart

## Abstract

SARS-CoV-2 virus has continued to evolve over time necessitating the adaptation of vaccines to maintain efficacy. Monoclonal antibodies (mAbs) against SARS-CoV-2 were a key line of defense for unvaccinated or immunocompromised individuals. However, these mAbs are now ineffective against current SARS-CoV-2 variants. Here, we tested three aspects of αSARS-CoV-2 therapeutics. First, we tested whether Fc engagement is necessary for *in vivo* clearance of SARS-CoV-2. Secondly, we tested bi-specific killer engagers (BiKEs) that simultaneously engage SARS-CoV-2 and a specific Fc receptor. Benefits of these engagers include the ease of manufacturing, stability, more cell-specific targeting, and high affinity binding to Fc receptors. Using both mAbs and BiKEs, we found that both neutralization and Fc receptor engagement were necessary for effective SARS-CoV-2 clearance. Thirdly, due to ACE2 being necessary for viral entry, ACE2 will maintain binding to SARS-CoV-2 despite viral evolution. Therefore, we used an ACE2 decoy Fc-fusion or BiKE, instead of an anti-SARS-CoV-2 antibody sequence, as a potential therapeutic that would withstand viral evolution. We found that the ACE2 decoy approach also required Fc receptor engagement and, unlike traditional neutralizing antibodies against specific variants, enabled the clearance of two distinct SARS-CoV-2 variants. These data show the importance of Fc engagement for mAbs, the utility of BiKEs as therapies for infectious disease, and the *in vivo* effectiveness of the ACE2 decoy approach. With further studies, we predict combining neutralization, the cellular response, and this ACE2 decoy approach will benefit individuals with ineffective antibody levels.

**Abbreviations:** ACE2, scFv, mAb, BiKE, COVID-19, Fc, CD16, CD32b, CD64, d.p.i

**Key points:**

- With equal dosing, both neutralization and Fc engagement are necessary for the optimal efficacy of *in vivo* antibodies and bi-specific killer engagers (BiKEs) against SARS-CoV-2.
- BiKEs can clear SARS-CoV-2 virus and protect against severe infection in the hACE2-K18 mouse model.
- ACE2 decoys as part of Fc-fusions or BiKEs provide *in vivo* clearance of two disparate SARS-CoV-2 variants.

## Introduction

Coronavirus disease 2019 (COVID-19) is caused by severe acute respiratory syndrome coronavirus 2 (SARS-CoV-2) and has led to 778 million deaths since it was first identified in 2019^1^. Over the past four years, there have been five variants of concern (VOCs), including Alpha, Beta, Gamma, Delta and Omicron, that have caused multiple waves of infection^2^. These variants have consistently accumulated mutations in the Spike protein of SARS-CoV-2^3^. The Spike protein binds angiotensin-converting enzyme 2 (ACE2) receptors on the surface of human cells to enter them and so is the immunization target for SARS-CoV-2. The mutations in Spike have resulted in increased transmissibility and resistance to infection- or vaccine-induced antibodies^4–8^. For example, antibodies that were previously shown to be capable of neutralizing the original Alpha variants are ineffective against the latest Omicron variants^6^. Reinfection by neutralization-escape variants and break-through infections in those that have been vaccinated is now commonplace against SARS-CoV-2 infection^2,9,10^. Vaccine companies will continue to incorporate new variants of SARS-CoV-2 into their vaccines. However, there is still an urgent need for therapeutics for people who cannot generate effective antibody levels, or who may have an adverse reaction to vaccine components. These individuals include immunocompromised patients, like leukemia patients, that had decreased morbidity and death with anti-SARS-CoV-2 mAb therapy^11,12^. Antiviral drugs partially address this issue, however, effective monoclonal antibody therapies that can withstand viral evolution would significantly benefit these patient populations. Furthermore, different viruses have unique protective and pathological antibody responses, such as influenza and Dengue virus respectively^13–15^. For example, in Dengue, antibodies can interact with Fc receptors to cause antibody-dependent enhancement in which certain concentrations of antibodies do not prevent infection, but rather exacerbate it^14^. Therefore, understanding which Fc-mediated antibody responses are beneficial and do not cause adverse effects is crucial information for SARS-CoV-2.

Antibodies have multiple functions, and many factors can determine whether a mAb will be effective or not. Understanding what makes antibodies effective at clearing virus is necessary to develop therapeutics that will continue to work even with SARS-CoV-2 evolution. One factor to consider is whether to target the neutralizing or non-neutralizing surface targets of the pathogen. Neutralization is defined by the antibody blocking the virus from being able to enter and infect a permissive cell type. In the case of SARS-CoV-2, a neutralizing antibody can block the ability of the receptor binding domain (RBD) of the Spike protein to bind to ACE2 or prevent the cleavage of Spike for the next stage of viral entry^16,17^. A non-neutralizing antibody is one that binds the surface of the pathogen but does not block entry into permissive cells, such as the non-RBD/cleavage portions of the Spike, Membrane, or Envelope proteins exposed on the SARS-CoV-2 viral surface^3^. Both neutralizing and non-neutralizing antibodies can bind to Fc receptors on effector cell types via their Fc binding portion^18–21^. Subsequently, a cytotoxic cell such as a natural killer (NK) cell can initiate destruction of the virus budding from, or on the surface of, a virus producing cell via antibody-dependent cellular cytotoxicity (ADCC).

Alternatively, a phagocytic cell can engulf the ‘free’ extracellular virus or the virus on the surface of an infected cell through antibody dependent cellular phagocytosis (ADCP)^22^. Much work has been done in comparing neutralizing and non-neutralizing monoclonal antibodies in influenza and HIV research^15,19,23–25^. The consensus from that work is that neutralizing monoclonal antibodies that can bind Fc receptors on effector cells are more effective than non-neutralizing antibodies when injected at similar quantities. However, at higher concentration, some non-neutralizing antibodies can be effective at clearing virus^15,18,24^. With SARS-CoV-2, the role of Fc effector functions in protection against COVID-19 disease is not fully clear. In humans, Fc-mediated effector function activity is generally connected with neutralization activity as wildtype antibodies both neutralize and have Fc domains, making it difficult to delineate which antibody function (neutralization or Fc-mediated effector functions) affects disease outcome^26^. For mechanistic and therapeutic purposes, mutations have been identified in the antibody Fc portion that ablate binding to Fc receptors that can help delineate these functions^27^. With current tools and considering previous data, we wanted to determine what characteristics, including neutralization and Fc receptor engagement, are most effective for therapeutics targeting of SARS-CoV-2.

In addition to improving currently available canonical mAbs, there is also interest in exploring ‘killer engagers,’ which are engineered hybrid, typically bispecific molecules, designed for more efficient and targeted killing of cancer cells or pathogens. One side is an antibody derivative (scFv or nanobody) that is specific for the cancer or infectious disease target and the other side is an antibody-like component that is specific for a particular Fc receptor or the T cell receptor. These killer engagers have previously been used to increase the killing activity of NK cells (bi-specific killer engagers or BiKEs) or T cells (bi-specific T cell engagers or BiTES) in the context of cancer immunotherapy^28–30^. Furthermore, these bispecific molecules can be improved by incorporating other signaling molecules. For example, for NK cells, recombinant IL-15 can be added to the BiKE, now called tri-specific killer engagers (TriKEs), to further improve NK cell function. Current TriKEs are aimed to bind cancer cells, activate NK cells through IL-15, and engage in ADCC through their CD16a (FcγRIIIa) Fc receptor^28,29^. There are many advantages to using ‘killer engagers’ over monoclonal antibodies^29,31^: 1) terminal end nanobody proteins are a single amino acid chain that are easier to manufacture and more thermally stable; 2) they target certain Fc receptors with higher affinity than antibodies (antibodies have variable affinity for Fc receptors because of different antibody glycosylation patterns and host Fc receptor genetics that affect IgG Fc receptor binding affinity); 3) antibody isotypes bind multiple Fc receptors so using specific anti-Fc killer engagers enables targeting of certain cytotoxic or phagocytic cell types more specifically; 4) IgG antibodies also bind the inhibitory Fc receptor FcγRIIb (CD32b) which has been shown to cause antibody dependent enhancement (ADE) in other viral infections like Dengue virus. The utility of killer engagers in infectious diseases is unknown.

As we have seen mAbs become ineffective against new SARS-CoV-2 variants, there is a strong need to identify therapeutics that are able to neutralize SARS-CoV-2 as its sequence evolves. Many studies have suggested the use of ACE2 protein as a decoy to maintain efficacy against multiple SARS-CoV-2 variants. This is an alternative strategy to broadly neutralizing antibodies against SARS-CoV-2. ACE2, the protein SARS-CoV-2 binds to for viral entry, can therefore be used as a soluble decoy receptor that competes for binding to the Spike protein^32–48^. In principle, if a Fc-fusion or BiKE has an ACE2 decoy on one end, once it binds to SARS-CoV-2, the ACE2 decoy will neutralize the virus and effector cells will recognize the Fc-fusion/BiKE and clear the bound virus or kill the virus producing cell. The advantage of the ACE2 decoy is that it is resistant to SARS-CoV-2 mutations because SARS-CoV-2 cannot mutate to the extent that it loses binding to ACE2 as this would halt its life cycle. In previous studies, engineered higher Spike affinity ACE2 decoy Fc fusions successfully improve blockade and neutralize multiple viral variants of SARS-CoV-2 *in vitro* and *in vivo*^32,35,37,39,40,42–44,46,49–52^. To our knowledge no study has assessed BiKEs employing the ACE2 decoy approach. Therefore, combining the ACE2 decoy and BiKE machinery may ultimately maintain neutralization during viral evolution, which increases options to target different cellular effectors for injectable therapeutics and reduces the potential for ADE. Using the entry receptor as a decoy is being explored in HIV, where a modified CD4 domain binds HIV and can clear HIV infected cells when it is utilized as a chimeric antigen receptor (CAR)^53,54^. Further work is needed on the effectiveness of ACE2 decoy therapeutics *in vivo* against SARS-CoV-2 infection to know if they have promise in the clinic.

Thus, the overarching goal of this study was to provide initial data on the characteristics of effective injectable antibody-oriented therapeutics to treat individuals with ineffective levels of SARS-CoV-2 antibodies. We sought to test our hypothesis’ *in vivo* with multiple SARS-CoV-2 variants in K18-hACE2 mice, which express human ACE2^48,55,56^. We utilized known antibody systems to modulate Fc binding of antibodies by expressing antibodies that contain the loss of function LALA-PG substitutions in the Fc portion of the antibody, rendering the antibodies unable to bind to Fc γ receptors (∅FcγR)^27,57^. Furthermore, we utilized known neutralizing and non-neutralizing antibody sequences to assess the contribution of neutralizing and non-neutralizing antibodies and BiKEs in the protection from SARS-CoV-2^58^. Lastly, using sequences previously published, we engineered and tested high affinity (ha) ACE2 mutant decoy molecules that can or cannot bind Fc receptors^33,35,37,46,49,59^. It was not previously known if haACE2 Fc-fusions or novel haACE2 decoy BiKEs would be effective at clearing SARS-CoV-2 *in vivo* with or without Fc engagement. Our data highlights the synergy of both neutralization and cellular engagement for an effective injectable antibody/BiKE treatment. By utilizing both neutralization and the cell mediated response, we believe future work combining ACE2 decoys with ‘killer engagers’ may yield new tools for clinicians to treat individuals with ineffective antibody levels and that this treatment will endure SARS-CoV-2 evolution.

## Methods

### Mice

K18-hACE2 (Stock No. 034860) and C57BL/6J (B6) mice (Stock No. 000664) mice were purchased from The Jackson laboratory. FcγRI/II/III/IV quadruple KO (FcγRec1234^-/-^) mice were obtained from Dr. Sjef Verbeek^60^. All mice were propagated at the University of Minnesota. Mice were housed in groups of 3 to 5 per cage and maintained on a 12 h light/12 h dark cycle with access to water and standard chow diet *ad libitum*. All mice were grown in specific pathogen-free (SPF) condition. Both male and female mice between the ages of 8-16 weeks were used in this study.

### Maintenance of SARS-CoV-2 strains

Vero E6 cells (ATCC CRL-1586, Manassas, VA, USA) were grown in Dulbecco’s modified Eagle medium (DMEM) supplemented with 5% heat inactivated fetal bovine serum (FBS). SARS-CoV-2 isolate hCoV-19/South Africa/KRISP-EC-K005321/2020 (NR-54008; Lineage B.1.351), and isolate hCoV-19/USA/MD-HP47946/2023 (NR-59503; Lineage EG.5.1; Omicron Variant) were obtained from BEI Resources, NIAID, NIH (Manassas, VA, USA) and propagated in Vero E6 cells (ATCC CRL-1586) or Vero E6-TMPRSS2-T2A-ACE2 cells (BEI Resources NR-54970). Virus titers were determined by focus-forming assay on Vero E6 cells. All procedures with infectious SARS-CoV-2 were performed in certified biosafety level 3 (BSL3) facilities at the UMN using appropriate standard operating procedures (SOPs) and protective equipment approved by the UMN Institutional Biosafety Committee.

### Infection of Mice with SARS-CoV-2

K18-hACE mice in the infected group were anesthetized with 3% isoflurane/1.5 L/min oxygen in an induction chamber and inoculated intranasally with 1 x 10^4^ plaque forming unit (PFU) of SARS-CoV-2 in a volume of 50 μL DMEM, split equally between each nostril^55,56^. Mice were monitored, and the body weight measured daily for the duration of the experiment. At 5-6 days post infection (d.p.i), mice were euthanized, and lungs were harvested for downstream analysis.

B6 or FcγRec1234^-/-^ mice were anesthetized with 3% isoflurane/1.5 L/min oxygen in an induction chamber and inoculated intranasally with 1 x 10^9^ PFU adeno-associated virus (AAV) vector that transiently expresses human ACE2 in the nasal and lung passages^50^. Five days later, they were anesthetized again with 3% isoflurane/1.5 L/min oxygen in an induction chamber and inoculated intranasally with 1 x 10^6^ PFU of SARS-CoV-2 in a volume of 50 μL DMEM, split equally between each nostril. Mice were monitored, and the body weight measured daily for the duration of the experiment. At 6 days d.p.i, mice were euthanized, and lungs were harvested for downstream analysis.

### Generation of SARS-CoV-2 binding antibodies, Fc-fusion proteins, and BiKEs

The amino acid (AA) sequences of the neutralizing (CC12.1), and non-neutralizing (CC12.21) SARS-CoV-2 binding variable heavy (VH) and variable light (VL) antibody domains were first described in Rogers et al^58^. To generate SARS-CoV-2 neutralizing antibodies with or without Fc receptor binding (∅FcγR), neutralizing (CC12.1) VH and VL were cloned into previously described pcDNA3.4 expression vectors containing either mIgG2a or mIgG2a-LALA-PG, and mIgK antibody constant domains, respectively^61^. The LALA-PG contains three point mutations in the Fc binding portion that render the Fc portion of the antibody null (∅FcγR): L234A, L235A, P329G^62^. Cloning was performed via Gibson assembly, with mouse codon optimized gene fragments for the VH and VL domain containing 30 base pair 5’ and 3’ homologous overlaps to the linearized destination antibody heavy and light chain expression vector. Binding (or lack thereof) to Fc receptors was confirmed using Bio-Layer Interferometry (Octet) (Supplemental Table 1).

The AA sequence of the Spike high affinity human ACE2 variant 118 (haACE2), was first described in Glasgow, et al^49^. To generate SARS-CoV-2 neutralizing hACE2_118_ Fc-fusion proteins with or without Fc receptor binding (∅FcγR), hACE2_118_ (encompassing AA residues 18 to 614, and thus lacking the collectrin domain) was cloned into the aforementioned mIgG2a and mIgG2a-LALA-PG heavy chain expression vectors via Gibson assembly, as above, with the mIgG2a and mIgG2a-LALA-PG expression vectors were linearized with AflII and PspOMI. PspOMI cuts within the coding sequence of the mIgG2a hinge domain such that the translated hinge region would be modified to remove only a Pro and Arg residue at the N-terminus of the hinge domain. Mouse codon optimized gene fragments were synthesized for haACE2, which included a Gly-Gly-Gly-Gly-Ser (G4S) linker before the slightly truncated hinge domain to fuse the haACE2 to the Fc domain of the antibody. The resulting linker between the hACE2_118_ domain and mIgG2a Fc domain was GGGGSGPTIKPCPPCKCPAPNLL

To generate the αSARS-CoV-2 BiKEs, sequences for single-chain variable fragments (scFvs) that bind to CD16, CD32b, and CD64 were obtained from previously described antibodies^63^ and VH and VL sequences of a null-binding control scFv, which we are calling “∅FcγR BiKE,” were obtained from the PDB (PDB ID: 1IGC) (classically called the ‘MOPC’ isotype control)^64,65^. Gene fragments encoding an scFv for the already described SARS-CoV-2 binding CC12.1 (neutralizing), and CC12.21 (non-neutralizing) antibody domains^58^, with the coding region from N-term to C-term oriented as VH-(G4S)_(3)_-VL, were synthesized and combined with gene fragments encoding the scFv of either anti-CD16, CD32, CD64, or ∅FcγR for Gibson assembly into a pcDNA3.4 expression vector downstream of the Kozak sequence and a serum murine albumin signal peptide (MKWVTFISLLFLFSSAYS). The two scFv of the BiKE expression constructs were separated by a GSTSGSGKPGSGEGSTKG linker^66^ and followed by C-term 10x-Histidine tag. For haACE2 BiKE CD16, CD64, and ∅FcγR scFvs were connected by a GGGGSGSTSGSGKPGSGEGSTKG linker and C-term 10x-Histidine tag ^49^.

All gene fragments were ordered from Twist Biosciences. Recombinant protein expression was performed with the Expi293 expression system (ThermoFisherScientific, Catalog # A1463) according to the manufacturer’s instructions. All proteins were purified with an AKTA pure chromatography system (Cytiva). All Fc-containing proteins were purified from clarified and filtered cell culture supernatant with a HiTrap Protein G HP (Cytiva, Catalog # 29048581) or a HiTrap MabSelect PrismA protein A column (Cytiva, Catalog # 17549851). All BiKEs were purified from clarified and filtered cell culture supernatant with nickel-affinity HisTrap excel chromatography column (Cytiva, Catalog # 29048586). Purified samples were buffer exchanged into PBS, pH 7.4, and concentration was determined by absorbance at 280 nm on a Nanodrop (ThermoFisherScientific). Confirmation of expected molecular weight and analysis of the purity of the recombinant proteins was obtained from SDS-PAGE analysis under reducing and non-reducing conditions (Supplemental Figure 1).

### Measurement of viral burden

The left lung was weighed and homogenized in 1 mL of RPMI supplemented with 10% FBS in GentleMACS^™^ M tube using GentleMACS^™^ Dissociator (Miltenyi Biotec) with RNA 2.01 program setting. Tissue homogenates were stored in aliquots at −80 °C until use. Total RNA was extracted from 160 μL of tissue homogenate using the QIAamp Viral RNA Mini Kit (Qiagen) according to the manufacturer’s instructions. The RNAse-Free DNAse set was used to digest DNA during RNA purification according to the manufacturer’s instructions (Qiagen). The purity of RNA was assessed by calculating the ratio of OD_260_/OD_280_. 1 μg of total RNA was reverse transcribed using the Roche Transcriptor First Strand cDNA synthesis kit (Roche Life Sciences) according to manufacturer’s instructions.

The cDNA was amplified RT-qPCR using Fast SYBR Green master mix (Thermo Fisher scientific, Waltham, MA) in a CFX96 Touch Real-Time PCR Detection System (BioRad) using virus-specific nsp14 primers for nucleocapsid: 5’-TGGGGYTTTACRGGTAACCT-3’ and 5’-AACRCGCTTAACAAAGCACTC-3’.The following reaction conditions were used: 95 °C for 3 min followed by 40 cycles of 95 °C for 10 s, 65.5 °C for 30 s, and 72 °C for 30 s. The specificity of RT-qPCR was assessed by analyzing the melting curves of PCR products. Ct values were normalized to the GAPDH gene. The fold change was determined by comparing the average Ct values for the nucleocapsid gene compared to the GAPDH gene using 2^−ΔΔCt^ method.

### Statistical analysis

Statistical analyses were performed using Prism version 9.5.1 (GraphPad, La Jolla, CA). Analysis of weight change was determined by a two-way Analysis of Variance (ANOVA). A Kruskal-Wallis non-parametric one-way ANOVA with Dunn’s multiple comparison test was used to analyze viral load and relative gene expression of RT-qPCR data. A p-value of less than 0.05 was considered significant.

### Study approval

Animal studies were carried out in accordance with the recommendations in the Guide for the Care and Use of Laboratory Animals of the National Institutes of Health. The protocols were approved by the Institutional Animal Care and Use Committee (IACUC) and the Institutional Biosafety Committee (IBC) at the University of Minnesota.

## Results

### Monoclonal antibodies against SARS-CoV-2 require Fc effector functions for viral clearance

The passive transfer of neutralizing mAbs against SARS-CoV-2 has conferred protection in multiple pre-clinical mouse models^27,50,67,68^. However, in addition to neutralization, the *in vivo* antiviral efficacy can also be due to Fc-dependent functions that promote clearance of the virus and virally infected cells through the innate and adaptive immune system^18,21,22,24,69^. To determine the role of Fc-mediated effector functions, we engineered wildtype and non-FcγR binding (∅FcγR) versions of both neutralizing and non-neutralizing antibodies that target the Spike protein of SARS-CoV-2 (Figure 1A)^58^. The non-FcγR binding versions contain the loss of function LALA-PG substitutions (L234A, L235A, P329G) in the Fc portion of IgG2a backbone (Fc portion) of the antibody, which eliminates binding to FcγRs and C1q in both mice and humans (∅FcγR)^27,70^. We hypothesized that at similar doses Fc-mediated effector functions were necessary for protection against SARS-CoV-2 infection, regardless of whether the scFv region of the antibody was a neutralizing or non-neutralizing sequence. To study this, we inoculated K18-hCE2 mice with SARS-CoV-2 by the intranasal route and then delivered a single 200 μg dose of each antibody or combination of antibodies (Figure 1B). While passive transfer of the neutralizing αSpike IgG2a antibody decreased weight loss and viral load in the lungs, passive transfer of the neutralizing αSpike IgG2a ∅FcγR and the non-neutralizing antibodies with or without Fc function did not (Figure 1C). Combining a neutralizing αSpike IgG2a ∅FcγR antibody and a non-neutralizing αSpike IgG2a decreased viremia in the lung and reduced weight loss (Figure 1C). These data provide clear evidence that at similar doses of antibody, both neutralization and Fc mediated clearance synergize for effective viral clearance.

**Figure 1.**
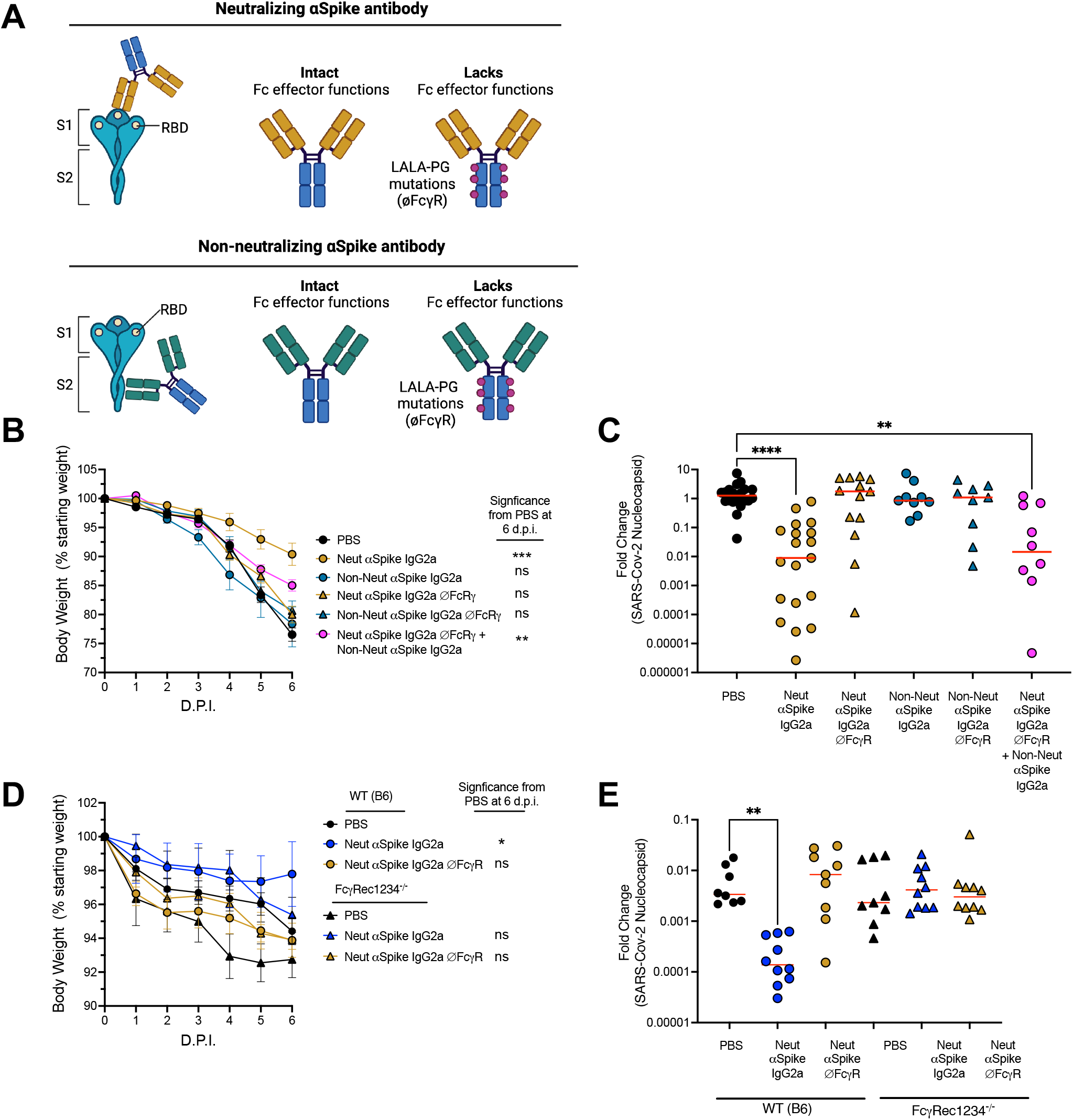
Fc effector functions increase the therapeutic activity of neutralizing antibody. **(A)** Schematic of neutralizing and non-neutralizing antibodies that are used. These antibodies are either SARS-CoV-2 neutralizing or non-neutralizing on the scFv portion of the antibody and either of the IgG2a isotype (IgG2a) or the IgG2a isotype modified with a LALA-PG mutation (IgG2a ∅FcγR) that cannot bind Fc γ receptors in mice. **(B)** 8- to-16-week-old K18-hACE2 mice were inoculated by intranasal route with 10^4^ PFU of the South African variant of SARS-CoV-2. At 1 d.p.i mice were given 200μg of neutralizing αSpike IgG2a, neutralizing αSpike IgG2a ∅FcRγ, non-neutralizing αSpike IgG2a, non-neutralizing αSpike IgG2a ∅FcγR, or a combination of these. Weight change in K18-hACE2 mice. Statistical analysis was only performed 6 d.p.i when all mice were alive to avoid survivor bias (mean +/- SEM, n=8, 3 experiments). **(C)** Viral load in the lung as quantified at 6 d.p.i. RT-qPCR for SARS-CoV-2 nucleocapsid RNA fold change comparing PBS to different mAb treatments. Red lines indicate median. N=18 per group, 4 experiments. A Kruskal-Wallis test with Dunn’s multiple comparisons was used to compare the mAb treat-groups to the PBS control at 6 d.p.i. *p<0.05, **p<0.005. **(D)** 8- to-16-week-old B6 or FcRγRec124^-/-^ mice were infected with 10^9^ PFU adenovirus vector that transiently expresses hACE2 in the nasal and lung passages. These mice were then inoculated by intranasal route with 10^6^ PFU of the South African variant of SARS-CoV-2. At 1 d.p.i mice were given 200μg of neutralizing αSpike IgG2a or neutralizing αSpike IgG2a ∅FcγR. Weight change in either B6 or FcRγRec1234^-/-^ mice. Statistical analysis was only performed 6 d.p.i when all mice were alive to avoid survivor bias (mean +/- SEM, n=18, 4 experiments). **(E)** Viral load in the lung as quantified at 6 d.p.i. RT-qPCR for SARS-CoV-2 nucleocapsid RNA fold change comparing nucleocapsid to GAPDH. Red lines indicate median. N=8 per group, 3 experiments. A Kruskal-Wallis test with Dunn’s multiple comparisons was used to compare the mAb treat-groups to the PBS control at 6 d.p.i. *p<0.05, **p<0.005.

As described above, the LALA-PG substitutions eliminate binding to FcγRs. However, to investigate this result with a different approach, we performed a similar experiment with a neutralizing IgG2a or neutralizing IgG2a with LALA-PG mutation (neutralizing or non-neutralizing IgG2a ∅FcγR) in wildtype B6 and FcγR1234^-/-^ mice, lacking all Fc γ receptors (Figure 1D). To infect B6 and FcγR1234^-/-^ mice that do not express human ACE2, we used an AAV system that transiently transfects the nasal and lung tissue with human ACE2 and then followed this with SARS-COV-2 infection. The only statistically significant effect was seen with the neutralizing IgG2a antibody in B6 mice (with Fc γ receptors). There were no differences between PBS-treated or ∅FcγR mAb-treated B6 animals. There were also no differences between mAb-treated and PBS-treated FcγRec1234^-/-^ mice (Figure 1E). These data provide evidence from two approaches that the Fc γ receptor interactions are needed for effective viral clearance.

### BiKEs that bind CD16 or CD64 and neutralize SARS-CoV-2 have therapeutic efficacy in K18-hACE2 mouse models

Next, we generated BiKEs where one side had the same neutralizing or non-neutralizing scFv sequences as in the experiments above and the other side contained αCD16 (FcγRIII), αCD64 (FcγRI), αCD32b (FcγRIIb), or non-binding control scFv (∅FcRγ) (Figure 2A, Supplemental Figure 2A). CD16 and CD64 are activating Fc γ receptors while CD32b is an inhibitory Fc γ receptor. We hypothesized that CD64 and CD16 would have a positive effect, while the inhibitory Fc receptor, CD32b, would lead to antibody-dependent enhancement, leading to a negative effect on viremia. A benefit of these BiKEs is the ability to target activating Fc γ receptors only, whereas antibodies, like human IgG1 that is current used in most antibodies used to treat SARS-CoV-2, bind both activating and inhibitory Fc γ receptors^11,12^. We found that a BiKE that both neutralized and targeted activating Fc γ receptors (neutralizing αSpike-αCD16 or neutralizing αSpike-αCD64) decreased weight loss and viremia.

**Figure 2.**
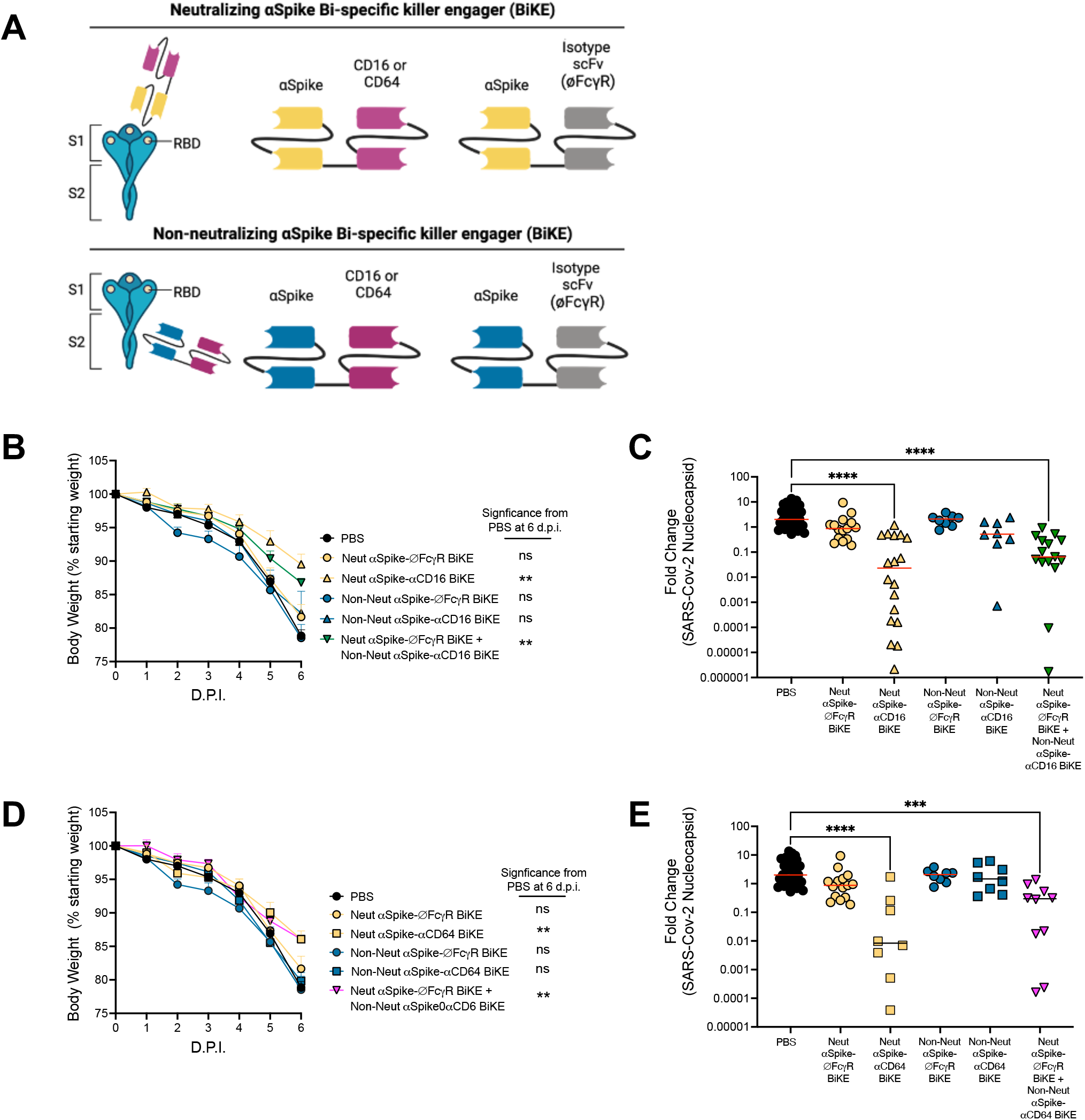
Fc effector functions increase the therapeutic activity of αSpike bi-specific killer engagers (BiKEs). **(A)** Schematic of neutralizing and non-neutralizing BiKEs that are used. On one end, these BiKEs have either a neutralizing scFv or non-neutralizing scFv. One the other end is an scFv against CD16 (FcγRIII), CD64 (FcγRI) or a MOPC null-binding isotype control scFv that binds to nothing in the mouse (∅FcγR). **(B-E)** 8-to-16-week-old K18-hACE2 mice were inoculated by intranasal route with 10^4^ PFU of the South African variant of SARS-CoV-2. At 1 d.p.i mice were given 50μg of the appropriate or combination of BiKEs. **(B,D)** Weight change. Statistical analysis was only performed 6 d.p.i when all mice were alive to avoid survivor bias (mean +/- SEM, n=8-15, 3 experiments). A Kruskal-Wallis test with Dunn’s multiple comparisons was used to compare the BiKE treated groups to the PBS control a 6 d.p.i. *p<0.05, **p<0.005. **(C, E)** Viral load in the lung as quantified at 6 d.p.i. RT-qPCR for SARS-CoV-2 nucleocapsid RNA fold change comparing PBS to different BiKE treatment treatments. Red lines indicate median. N=8-15 per group, 3 experiments. A Kruskal-Wallis test with Dunn’s multiple comparisons was used to compare the BiKE treated groups to the PBS control. ***p<0.005, ****p<0.00005

However, a BiKE that neutralized but did not bind to Fc γ receptors (neutralizing-∅FcγR) had little effect on weight loss or viremia in the lung (Figure 2B-E). If we injected non-neutralizing BiKEs that targeted Spike protein and Fc γ receptors CD16, CD64 or CD32b (non-neutralizing αSpike-αCD16, non-Neutralizing αSpike-αCD64 or non-neutralizing αSpike-αCD32b) there was no significant effect on weight loss or viremia (Figure 2B-E, Supplemental Figure 2). No evidence of ADE was seen in this system when targeting CD32b (Supplemental Figure 2). Lastly, if we injected both a neutralizing BiKE with no Fc γ receptor binding (neutralizing αSpike-∅FcγR) with a BiKE that was non-neutralizing with ability to bind Fc γ receptors (non-neutralizing αSpike-αCD16 or non-neutralizing αSpike-αCD64), we found decreases in viremia and reduced weight loss (Figure 2B-E). These data mirror those in Figure 1. These experiments again demonstrate that both neutralization and Fc γ receptor mediated cellular engagement is needed for effective viral clearance. It also shows that BiKEs to two different activating Fc γ receptors are capable of decreasing viremia *in vivo.* This to our knowledge is the first time this has been shown for BiKEs in an infectious disease.

### ACE2 decoy receptor Fc-fusions require Fc effector functions for viral clearance

ACE2 decoy receptors are an alternative strategy to anti-SARS-CoV-2 mAbs that bind and block the Spike protein^36,51^. We engineered high affinity (ha) ACE2 decoy receptor IgG2a Fc-fusion molecules with or without Fc receptor binding (∅FcγR) (Figure 3A). Like the αSpike antibodies in Figure 1, the non-Fc γ receptor binding versions contained the loss of function LALA-PG substitutions in the IgG2a backbone (Fc portion) of the antibody. We hypothesized that the haACE2 decoy Fc-fusion that can bind to Fc γ receptors would be the only version able to protect against SARS-CoV-2 infection. Of note, we had to increase the μg dose of the haACE2 Fc-fusion to achieve the molar equivalent dose of the neutralizing αSpike antibody (the haACE2 Fc-fusion is approximately double the molecular weight of the neutralizing antibody and BiKEs). Then, to see a significant effect, we found that we had to give 3 times the dose of the αSpike IgG2a mAb. We found that passive transfer of the haACE2 IgG2a Fc-fusion decreased weight loss and viral load in the lungs as well as the neutralizing αSpike IgG2a, while passive transfer of the neutralizing hACE2 IgG2a ∅FcγR Fc-fusion that did not (Figure 3B-C). Therefore, we conclude that a haACE2 Fc-fusion decreases viral load during SARS-CoV-2 infection and that it requires Fc γ receptor engagement for optimal function.

**Figure 3.**
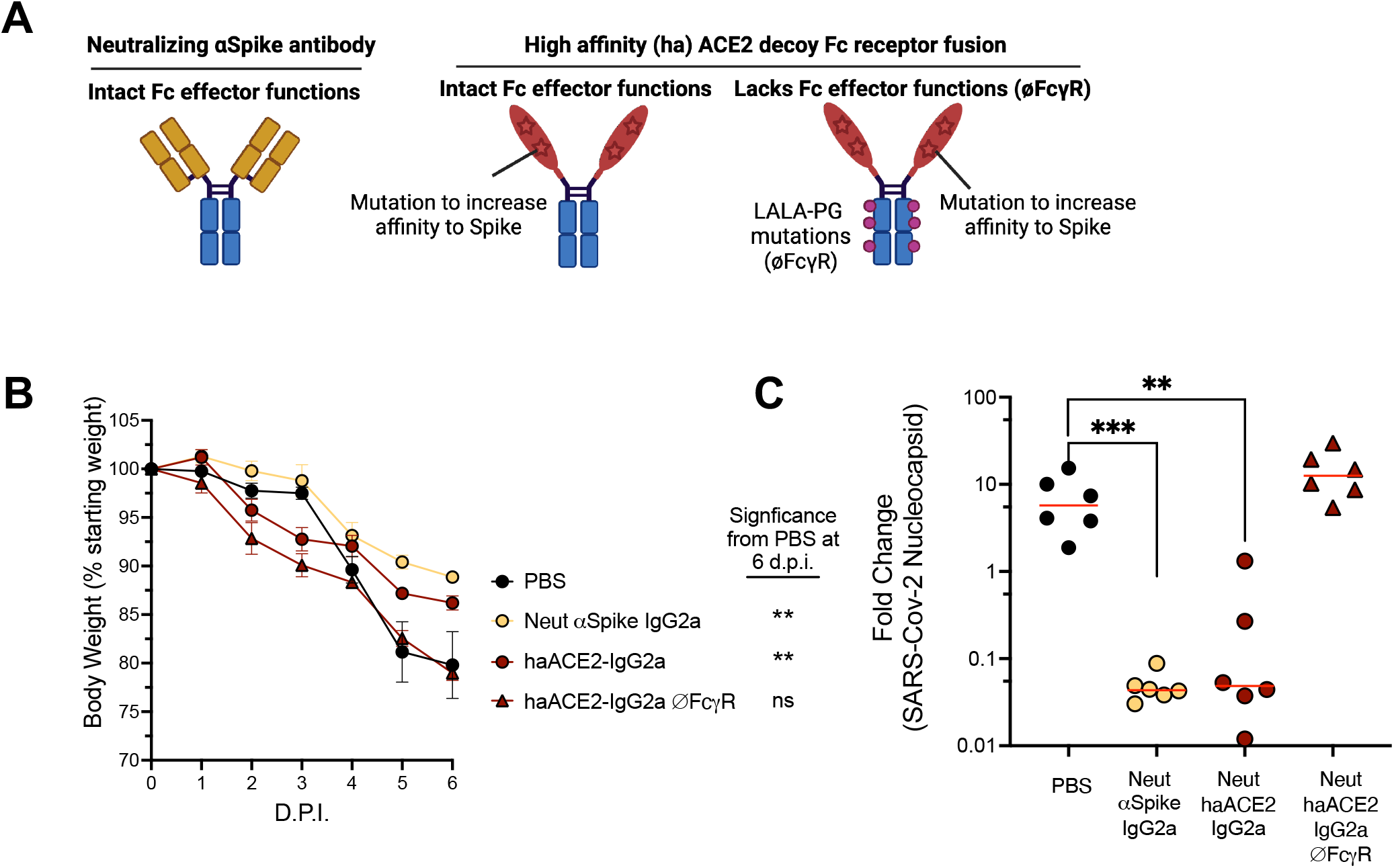
Fc effector functions increase the therapeutic activity of haACE2 decoy Fc-fusion. **(A)** Schematic of neutralizing and non-neutralizing antibodies that are used. One antibody is a SARS-CoV-2 neutralizing IgG2a antibody. The other antibodies, instead of a SARS-CoV-2 neutralizing scFv, containing a haACE decoy Fc-fusion. The ACE2 decoy receptors are either of the IgG2a isotype (IgG2a) or the IgG2a isotype modified with a LALA-PG mutation that cannot bind Fc receptors in mice (IgG2a ∅FcγR).**(B-C)** 8-to-16-week-old K18-hACE2 mice were inoculated by intranasal route with 10^4^ PFU of the South African variant of SARS-CoV-2. At 1 d.p.i mice were given 200μg of neutralizing αSpike IgG2a or at 1 d.p.i and 2 d.p.i were given 300μg haACE2 decoy IgG2a or hACE2 decoy IgG2a ∅FcγR. **(B)** Weight change in K18-hACE2 mice. Statistical analysis was only performed 6 d.p.i when all mice were alive to avoid survivor bias (mean +/- SEM, n=6, 2 experiments). A Kruskal-Wallis test with Dunn’s multiple comparisons was used to compare the mAb treat-groups to the PBS control a 6 d.p.i. **p<0.005. **(B)** Viral load in the lung as quantified at 6 d.p.i. RT-qPCR for SARS-CoV-2 nucleocapsid RNA fold change comparing PBS to different mAb treatments. Red lines indicate median. N=6 per group, 2 experiments. A Kruskal-Wallis test with Dunn’s multiple comparisons was used to compare the mAb treated groups to the PBS control. **p<0.005, ***p<0.0005

### BiKEs with an haACE2 decoy receptor on one side and an anti-Fc γ receptor scFv on the other side decreases viral load in both South African and Omicron SARS-CoV-2 variants

In addition to Fc-fusions, we tested BiKEs that have one side of the molecule that is the same haACE2 decoy used in Figure 3 and the other side is an scFv that binds to CD64 or ∅FcγR scFv (null-binding control) (Figure 4A). We hypothesized that the haACE2 decoy BiKEs would be effective against multiple variants of SARS-CoV-2 while the neutralizing αSpike BiKE would only be effective against the South African variant. This is because the sequence in the neutralizing αSpike BiKE generated against an earlier variant of the Spike protein (Washington variant); the Spike proteins of the South African and Washington strain are much closer in sequence than the Washington and Omicron variants. Like the haACE2 Fc-fusion, we had to increase the dose of the haACE2 decoy BiKEs to have an equivalent *in vivo* effect to that of the neutralizing αSpike IgG2a mAb. The haACE2 decoy BiKEs were approximated double the molecular weight of the neutralizing αSpike IgG2a BiKE. However, we needed to give triple the molar dose (300µg vs 50µg) and add a second dose 24 hours later to see significant reductions in weight loss and a decrease in a viral load (Supplemental Figure 3). Once we optimized the dosing to the South African variant, we found that the haACE2-αCD64 BiKE decreased weight loss and viremia like that of the neutralizing αSpike-αCD64 BiKE (Figure 4B-C). However, if a haACE2 BiKE that did not bind to Fc γ receptors was injected (haACE2-∅FcγR ΒιKE), this had no significant effect on weight loss or viremia in the lung (Figure 4B-C). Importantly, the haACE2 decoy BiKE was effective at preventing weight loss and decreasing viral load in <u>both</u> South African and Omicron variants while the neutralizing αSpike-αCD64 BiKE prevented weight loss in both variants but only decreased viremia in the South African variant (Figure 4D-E). This is the first time that an haACE2 decoy BiKE approach has been shown to be effective at clearing multiple variants of SARS-CoV-2. These results indicate that harnessing both neutralization and the cellular immune response with the ACE2 decoy ‘killer engager’ approach could be a promising therapeutic strategy to overcome immune-evading SARS-CoV-2 variants for those individuals that cannot generate an effective antibody response to SARS-CoV-2.

**Figure 4.**
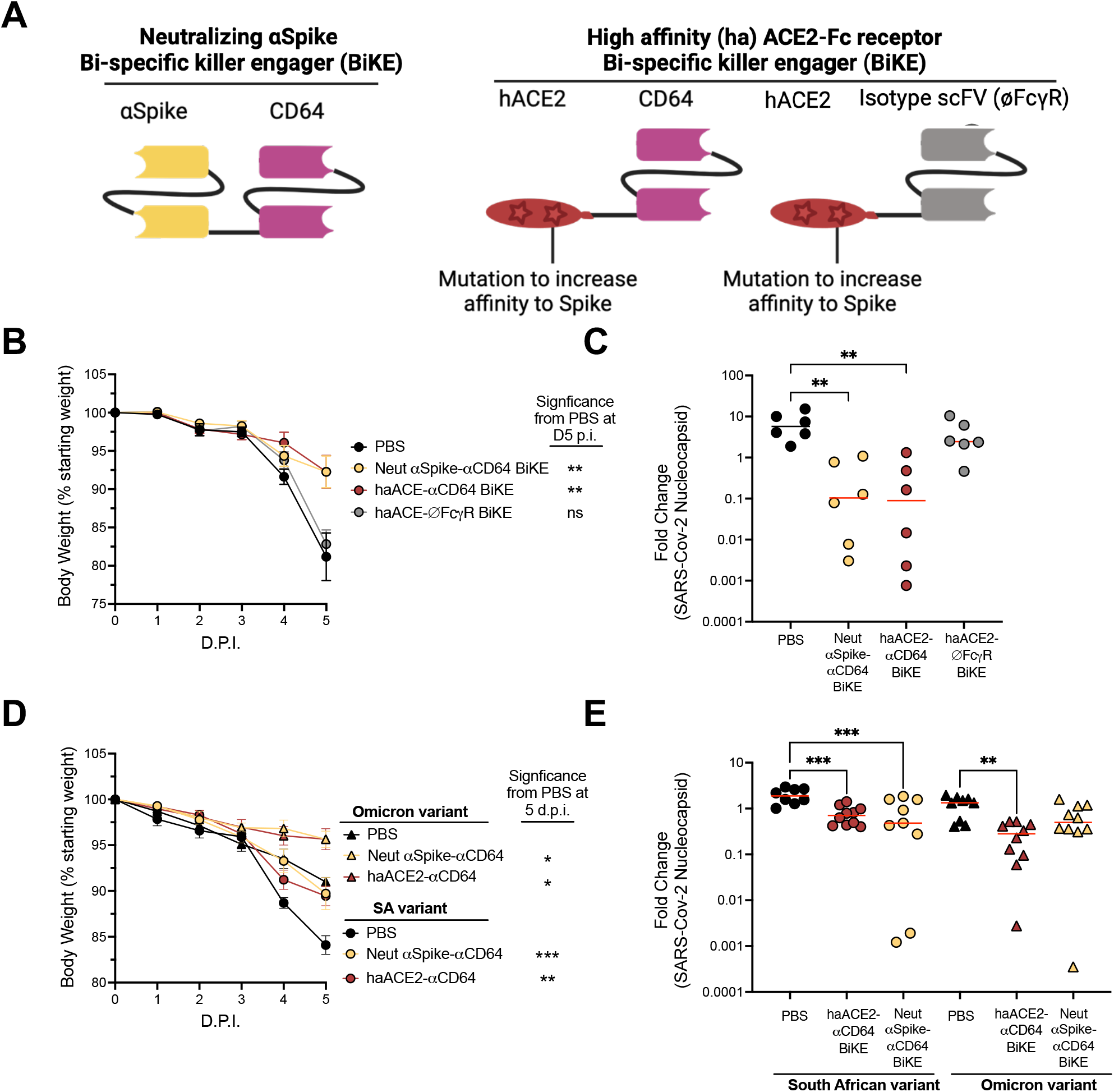
Fc effector functions increase the therapeutic activity of ACE2 decoy BiKE. **(A)** Schematic of neutralizing and non-neutralizing BiKEs that are used. On one end, these BiKEs have an ACE2 decoy receptor which contains mutations to increase affinity to the Spike protein that blocks the receptor binding domain. Fused to the ACE2 decoy receptor is an scFv against CD64 (FcγRI) binding or a MOPC null-binding control scFv (∅FcγR BiKE). **(B-E)** 8-to-16-week-old K18-hACE2 mice were inoculated by intranasal route with 10^4^ PFU of the South African or Omicron variant of SARS-CoV-2. At 1 d.p.i mice were given 50 μg of neutralizing αSpike CD64 BiKE or at 1 d.p.i and 2 d.p.i were given 300μg hACE2 decoy-CD64 BiKE or hACE2 decoy-∅FcRγ BiKE. **(B,D)** Weight change. Statistical analysis was only performed 6 d.p.i when all mice were alive to avoid survivor bias (mean +/- SEM, n=6-10, 2 experiments). A Kruskal-Wallis test with Dunn’s multiple comparisons was used to compare the BiKE-treated groups to the PBS control at 6 d.p.i. *p<0.05, **p<0.005, ***p<0.0005. **(C,E)** Viral load in the lung as quantified at 5 d.p.i. RT-qPCR for SARS-CoV-2 nucleocapsid RNA fold change comparing PBS to different BiKE treatments. Red lines indicate median. N=6-10 per group, 3 experiments. A Kruskal-Wallis test with Dunn’s multiple comparisons was used to compare the BiKE-treated groups to the PBS control. **p<0.005, ***p<0.0005

## Discussion

Novel therapeutics are needed that maintain efficacy amidst mutating SARS-CoV-2 variants to treat individuals that cannot develop their own effective antibody response. Therefore, it is necessary to understand what makes antibodies effective at clearing virus and to find approaches that will continue to work even with SARS-CoV-2 evolving. In this study, we show that the strongest therapeutic effect is delivered by antibodies that both neutralize Spike and bind Fc γ receptors on effector cells. We also show that effective clearance of SARS-CoV-2 was achieved with neutralizing BiKEs. Both approaches could be combined with a high affinity (ha) ACE2 decoy approach to effectively clear SARS-CoV-2 *in vivo*. Importantly, the haACE2 Fc-fusion and BiKE was effective at clearing two distinct SARS-CoV-2 variants (South African and Omicron), whereas a traditional neutralizing antibody was only effective for decreasing the viremia of the South African SARS-CoV-2 variant.

Our study further supports the findings that therapeutics that engage Fc γ receptors will be important for effective viral clearance^18^. At equivalent doses, lack of Fc γ receptor binding abolished any viral clearance seen by neutralizing antibodies and BiKEs. In agreement with our results, multiple studies have shown that SARS-CoV-2 neutralizing antibody protection is lessened by the loss of Fc-mediated functions in mouse models of COVID-19^27,71^. We also assessed the ability of non-neutralizing antibodies to protect against severe disease, with or without Fc effector functions. At the doses in our study, we showed that non-neutralizing antibodies themselves, with or without Fc effector functions, were not enough to decrease viremia or protect against weight loss. However, combining non-neutralizing antibodies that bound Fc γ receptors with neutralizing antibodies that did not bind Fc γ receptors were able to protect against disease. This ‘add-back’ experiment, further indicates that both the neutralization and Fc γ receptor-mediated clearance synergize for effective viral removal. Interestingly, a recent study reported that non-neutralizing mAbs can utilize Fc-mediated mechanisms to lower viral load and prevent lung damage due to SARS-CoV-2 infection^72^. We hypothesize that the differences with our data may relate to different antibody sequences and concentrations used. It has been shown in influenza virus infection that varying specificities and increasing the concentration of non-neutralizing antibodies can have a beneficial effect^13,15^. Therefore, it is likely that both an antibody response that neutralizes and utilizes Fc γ receptors or that induces a high level of non-neutralizing antibody can help the anti-viral response. Importantly, all the experiments reported in this paper used similar concentrations of neutralizing and non-neutralizing therapeutics to provide a direct comparison of effectiveness.

In addition to antibodies as therapeutics, we also explored the potential of using BiKEs to protect from SARS-CoV-2 infection. The potential of using BiKEs as therapeutics in SARS-CoV-2 was previously unknown. BiKEs and TriKEs have been utilized for more than a decade in the cancer setting^28,29^. A recent report showed that in Multisystem Inflammatory Syndrome in Children (MIS-C) BiKEs and TriKEs significantly increased the response of NK cells that were hyporesponsive due to Fc γ receptor genetics or patient group status^3^. This report also showed that monocytes that express CD64 were hyperactive and the anti-CD16 BiKE did not increase the antibody dependent cellular phagocytosis on monocytes, and only increased the NK cell activity in an *in vitro* PBMC assay. Therefore, unlike antibodies, these ‘killer engagers’ can be given individually or in combination to skew a high affinity interaction to one or multiple Fc γ receptors, while not engaging inhibitory Fc γ receptors, like FcγRIIb. Here, we created and tested BiKEs that targeted either CD16 or CD64. We showed that both Fc γ receptors can be targeted to protect against severe SARS-CoV-2 disease. In our K18-hACE2 mouse model, we saw a decrease in viremia over 6 days with the neutralizing anti-CD64 BiKE. Our experiment did not address long term persistence of virus or overall inflammation, but we did see decreased weight loss as an indirect measure of disease severity was reduced by treatment with BiKE.

One limitation of our model is that the K18-hACE2 mouse does not recapitulate physiological expression of ACE2. A main caveat of this mouse model is the abundant expression of human ACE2 throughout the animal; we believe this is not a major concern in this preliminary study as our main objectives were to decrease viremia and weight loss^48,55,56^. Further work using SARS-CoV-2 infections in more physiological hACE2 knock-in mice or primate models would address this caveat.

Furthermore, mouse CD16 is expressed by many cell types, including NK cells, myeloid cells, and neutrophils. CD64 expression is increased on mouse monocytes and on neutrophils after inflammation^22,69^. Unlike humans, mouse NK cells do not perform antibody dependent cellular cytotoxicity (ADCC) efficiently^73^. Therefore, we hypothesize that monocytes and neutrophils are the cellular effectors of protection with the CD64 and CD16 BiKEs. Conditional cell-specific Fc γ receptor ablation models would be needed to determine what proportion of the clearance was attributed to specific cell types.

There are reports that SARS-CoV-2 can infect lung resident macrophages and monocytes, and NK and CD8+ T cells can kill these infected cells in an IFNγ dependent manner^74–76^. If the macrophage infection phenotype is a potential risk for infection, this would further justify targeting specific Fcγ receptors versus using the canonical Fc of monoclonal antibodies that are more promiscuous binders to Fc γ receptors. For example, one could target specifically human CD16+ NK cells as opposed to CD64+ macrophages/monocytes. This is because both alveolar lung macrophages and monocytes are predominantly CD16 negative^77^.

One of the other Fc γ receptors that we tested was CD32b, an inhibitory Fc receptor. We were surprised to find no deleterious effect of targeting CD32b with a BiKE on viremia or weight loss. We hypothesized that targeting CD32b would increase the infectivity of SARS-CoV-2 into immune and non-immune cells that express FcγRIIb, like endothelial cells^78,79^. In other viruses, such as Dengue, antibodies can elicit ADE where sub-neutralizing antibodies against the virus particles opsonize the virus and allow entry via formation of immune complexes that interact with Fc γ receptor^14^. We did not see significant ADE here, as readout by increased viremia, but this could be relevant for broader application of BiKEs for infectious diseases. Our system gets around the potential for ADE by creating BiKEs that are specific to certain activating Fc γ receptors, as opposed to a mAb that would bind multiple Fc γ receptors including the inhibitory Fc γ receptor.

Lastly, based on our data showing synergy between neutralizing and Fc γ receptor binding approaches, we sought to find a way to maintain neutralization in the presence of an ever-evolving SARS-CoV-2 virus. Finding a broadly neutralizing antibody is one approach that has been tried with limited success^2,6,80,81^. We chose to explore the haACE2 decoy Fc-fusion approach because SARS-CoV-2 will need to maintain its binding to ACE2 to continue its life cycle, whereas this is not the case for any broadly neutralizing antibody. We found that an haACE2 decoy Fc-fusion and haACE2 decoy BiKE that engages an Fcγ receptor was successful at decreasing viremia and weight loss. However, we did have to increase the dose of the haACE2 Fc-fusion and BiKEs to have an equivalent *in vivo* effect to that of the neutralizing antibody and BiKEs. These doses are still well below the equivalent mg/kg dose given to humans, so this effective dose was within translational limits. We hypothesize this difference in dosing is due to the affinity difference between the neutralizing antibody and the haACE2 decoy Fc-fusion. For future experiments, we could increase the avidity of the haACE2 Fc-fusion by adding ACE2 molecules to help it stay bound to the virus. Additional work could also increase the affinity and neutralization potency of the haACE2 decoy Fc-fusion^33,49,82^. However, using an ACE2 decoy Fc-fusion that is as close to the human ACE2 sequence as possible while maintaining *in vivo* viral clearance efficacy, may give the longest lasting therapeutic effect to fight SARS-CoV-2 as it evolves. Alternatively, multiple ACE2 decoy Fc-fusions of higher affinity in different forms could be an effective long-term strategy. Importantly, as hypothesized, the haACE2 decoy approach maintained efficacy across multiple SARS-CoV-2 variants, that did not occur with the mAb.

In summary, we show that antibodies and BiKEs that are neutralizing and target Fc γ receptors decrease viremia and weight loss in the SARS-CoV-2 infected K18-hACE2 mouse model. Furthermore, we show that an ACE2 decoy Fc-fusion and BiKE can decrease viremia in disparate SARS-CoV-2 variants. There are many benefits of ‘killer engagers’ and, combined with this ACE2 decoy approach, may benefit individuals that do not have effective SARS-CoV-2 antibody responses. Further work will need to explore more physiological human and humanized models to discern what human Fcγ receptors and haACE2 decoy ‘killer engagers’ are best to target for maximal benefit and least detrimental effect. These data are a strong first step to demonstrate *in vivo* efficacy of these novel approaches for SARS-CoV-2 clearance.

## AUTHOR CONTRIBUTIONS

Conceptualization: G.T.H.

Methodology: J.K.D., D.H., V.D.K., C.B., M.C.C, M.P., G.T.H.

Investigation: J.K.D., V.D.K., J.A.S, B.T.Z.

Validation: J.K.D., D.H., G.T.H.

Formal Analysis: J.K.D., D.H., G.T.H.

Resources: D.H., J.S.V., M.S.C.

Data Curation: J.K.D., G.T.H.

Writing – Original Draft: J.K.D., G.T.H.

Writing – Review & Editing: All authors reviewed and edited the manuscript.

## Acknowledgements

This work was supported by the Department of Medicine, Department of Pharmacology, the Department of Veterinary Medicine, and the Clinical and Translational Science Institute at the University of Minnesota. This work was also supported by the National Institutes of Health grants ULTR1002494 and T32AI007313-34A1 (JKD). Thanks to the innovative work by Dr. Jeffrey Miller, Dr. Martin Felices, Todd Lenvik, and Dr. Yvette Soignier for the CD16 BiKE collaboration. We also thank Dr. Burton and Dr. Wells for publishing their sequences for their SARS-CoV-2 antibodies and ACE2 decoy, respectively. A special thanks to University of Minnesota BSL-3 program staff for their support of the high containment research laboratories used for these studies. Multiple figures made with Biorender.

## Model Figure

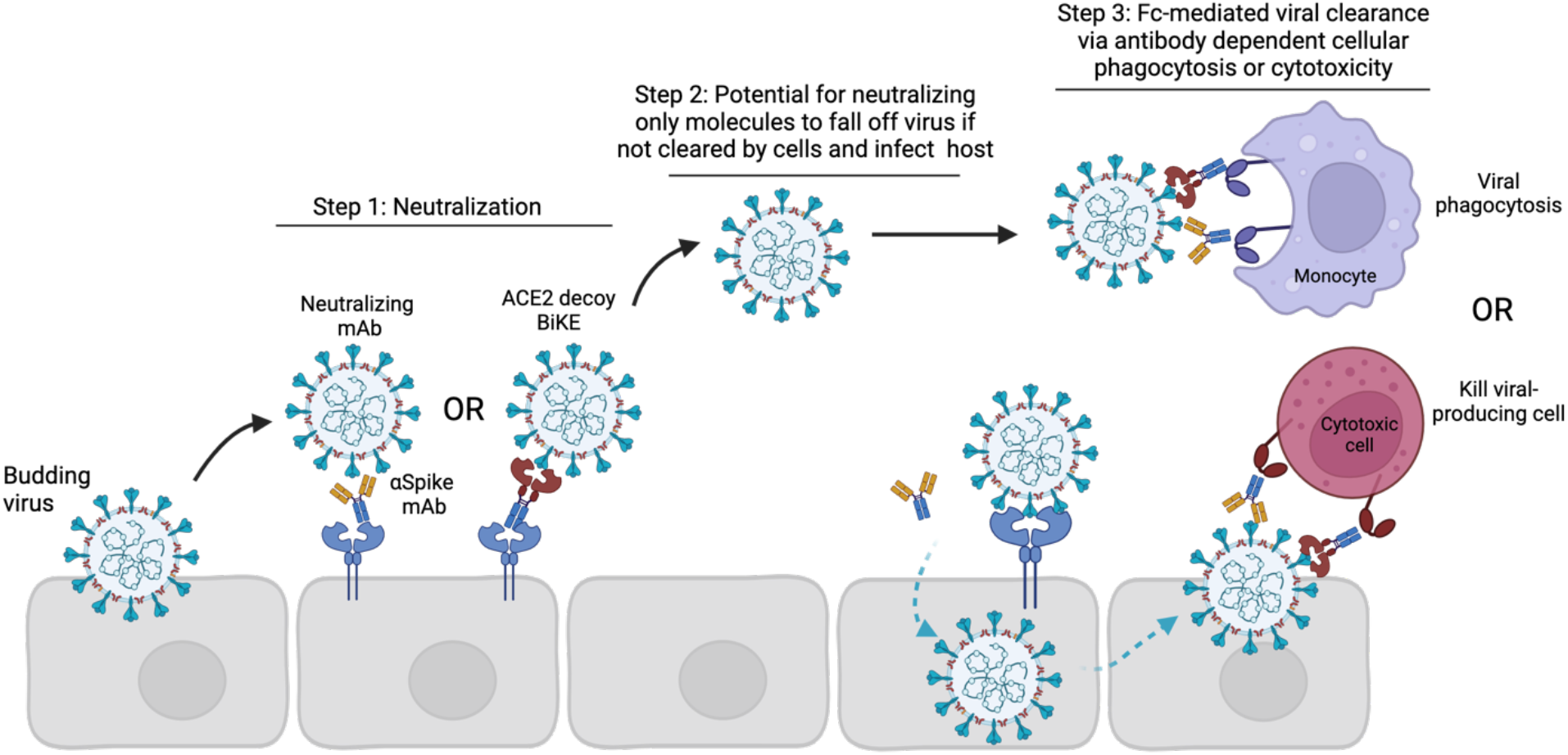

