## Supplemental Figures and Table for "ACE2 decoy Fc-fusions and bi-specific killer engager (BiKEs) require Fc engagement for *in vivo* efficacy against SARS-CoV-2"

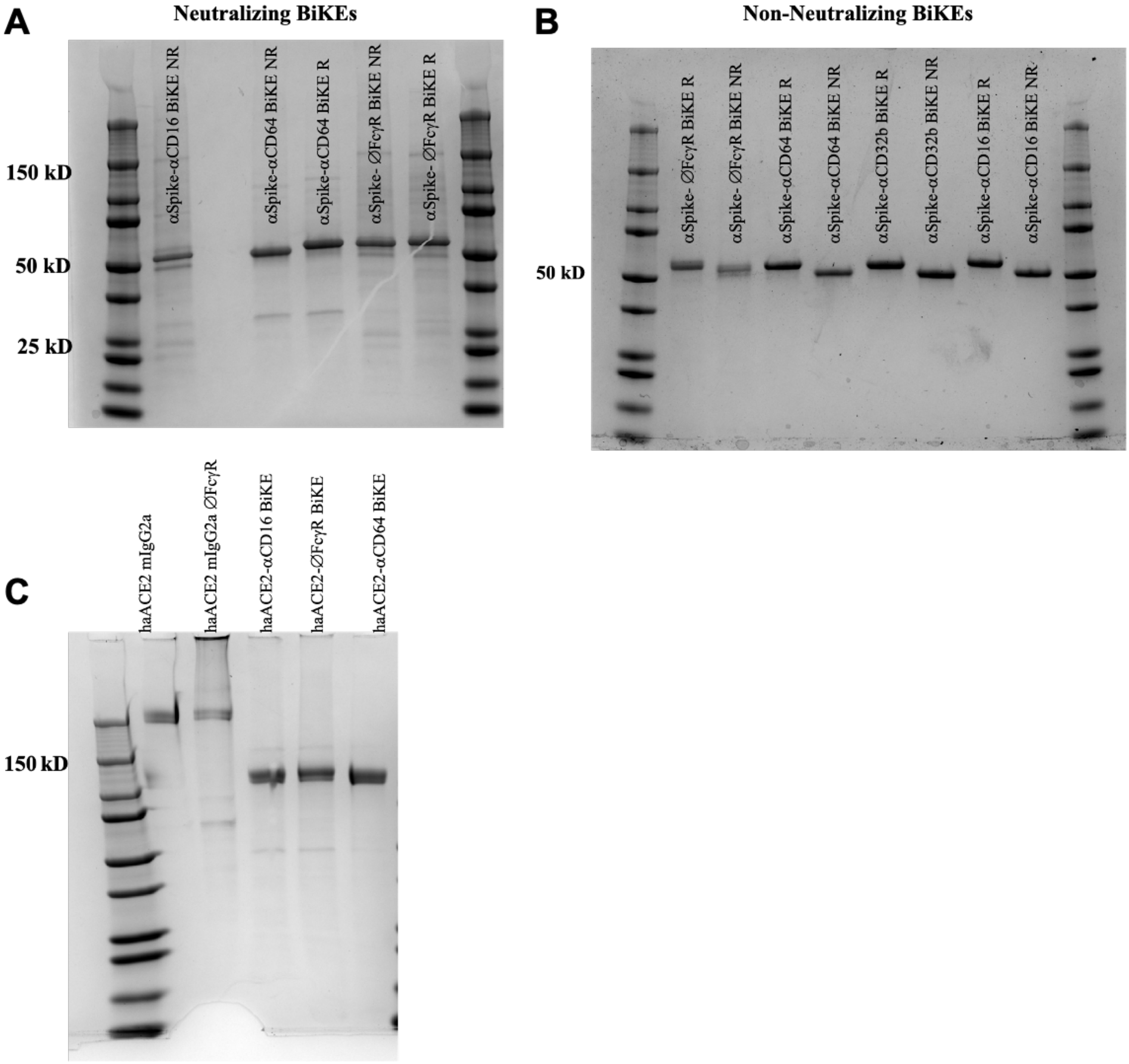


**Supplemental Figure 1.** Confirmation of expected molecular weight and analysis of the purity of the recombinant proteins. **(A)** SDS page gel of neutralizing BiKEs that are used. **(B)** SDS page gel of non-neutralizing BiKEs that are used. **(C)** SDS page gel of ACE2 decoy Fc-fusions that are used.

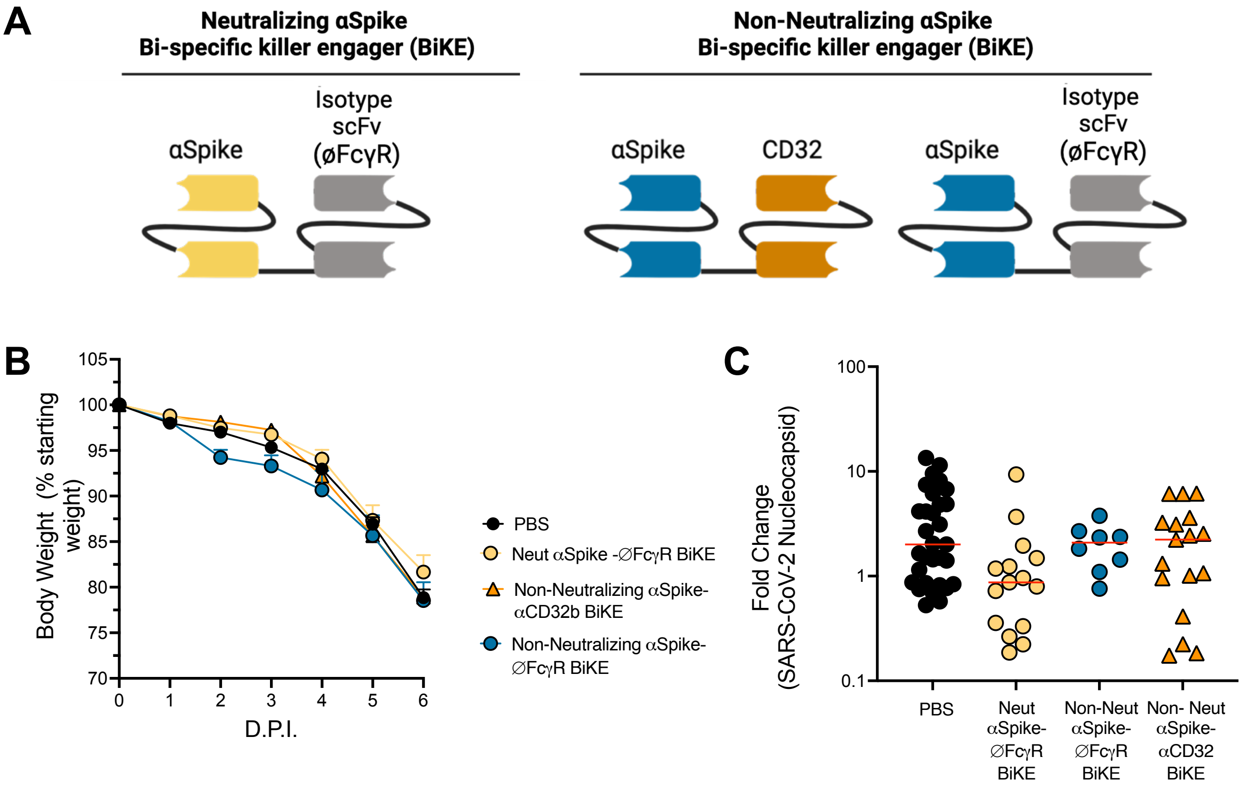


**Supplemental Figure 2. αSpike-CD32b BiKEs does not improve weight loss or viral load in K18-mice (A)** Schematic of neutralizing and non-neutralizing BiKEs that are used. On one end, these BiKEs have either a neutralizing scFv or non-neutralizing scFv. One the other end is CD32 (FcγRIIb) or a MOPC isotype control that binds to nothing in the mouse (∅FcγR). **(B-C)** 8- to-16-week-old K18-hACE2 mice were inoculated by intranasal route with $10^4$ PFU of the South African variant of SARS-CoV-2. At 1 d.p.i mice were given 50μg of the appropriate BiKE. **(B)** Weight change. Statistical analysis was only performed 6 d.p.i when all mice were alive to avoid survivor bias (mean +/- SEM, n=10-18, 3 experiments). A Kruskal-Wallis test with Dunn's multiple comparisons was used to compare the BiKE-treated groups to the PBS control a 6 d.p.i. **(C)** Viral load in the lung as quantified at 6 d.p.i. RT-qPCR for SARS-CoV-2 nucleocapsid RNA fold change comparing PBS to different BiKE treatment treatments. Red lines indicate median. N=8-15 per group, 3 experiments. A Kruskal-Wallis test with Dunn's multiple comparisons was used to compare the BiKE treated groups to the PBS control.

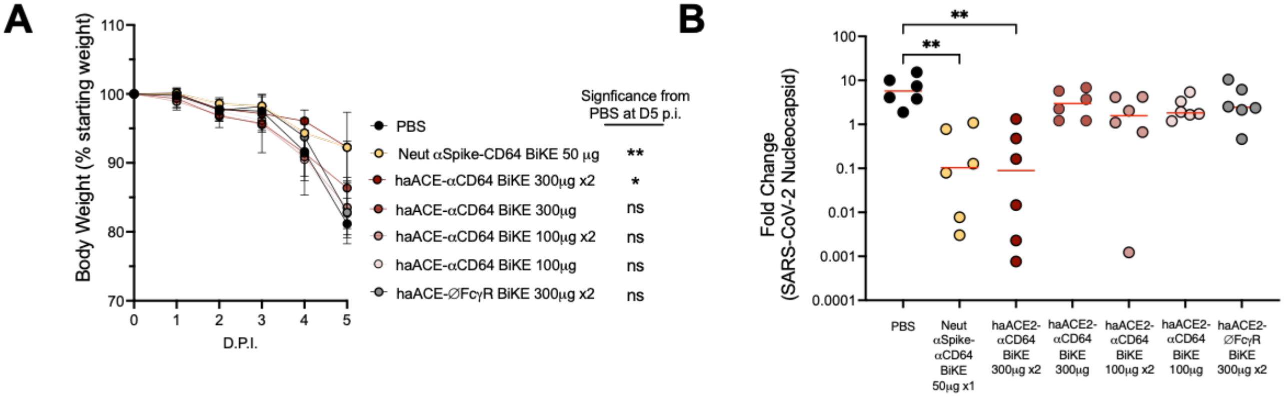


**Supplemental Figure 3. Dose escalation of ACE2 Fc fusion-CD64 BiKE shown *in vivo* viral load and weight loss treatment starting on Day 1 post infection. (A-B)** 12-week-old K18-hACE2 mice were inoculated by intranasal route with $10^4$ PFU of the South African or Omicron variant of SARS-CoV-2. Based on molecular weight 50µg of CD64-Spike is the same number of molecules as 100µg CD64-ACE. BiKEs were given once on day 1 post-infection ("x1") or given twice (once on day 1 and once on day 2 post-infection = "×2"). **(A)** Weight change. Statistical analysis was only performed 6 d.p.i when all mice were alive to avoid survivor bias (mean +/- SEM, n=6 per group, 2 experiments). A Kruskal-Wallis test with Dunn's multiple comparisons was used to compare the BiKE treated groups to the PBS control at 5 d.p.i. *p<0.05, **p<0.005. **(B)** Viral load in the lung as quantified at 5 d.p.i. RT-qPCR for SARS-CoV-2 nucleocapsid RNA fold change comparing PBS to different BiKE treatments. Red lines indicate median. N=6 per group, 2 experiments. A Kruskal-Wallis test with Dunn's multiple comparisons was used to compare the BiKE treated groups to the PBS control, **p<0.005.

**Supplemental Table 1. Binding affinities of αSARS-CoV-2 neutralizing antibodies with intact or lacking Fc γ receptor binding.**

| **mAb ID** | **Fc Receptor ID** | **Observed KD (M)** | **Published KD[83] (M)** |
|---|---|---|---|
| α-SARS-CoV-2 neu-mAb **mIgG2a** | mFcγRI | 2.03E-08 | 1.69E-08 |
| | mFcγRIIB | 2.12E-06 | 1.63E-06 |
| | mFcγRIII | 7.85E-07 | 9.45E-07 |
| | mFcγRIV | 2.65E-08 | 5.23E-08 |
| α-SARS-CoV-2 neu-mAb **mIgG2a LALA-PG** | mFcγRI | NDB | N/A |
| | mFcγRIIB | NDB | N/A |
| | mFcγRIII | NDB | N/A |
| | mFcγRIV | NDB | N/A |
